# Disentangling unity from diversity in developmental psychopathology

**DOI:** 10.1101/616177

**Authors:** Lorenna Sena Teixeira Mendes, Guilherme Vanoni Polanczyk, Luis Augusto Rohde, Giovanni Abrahão Salum

**Author notes:** Corresponding author: Lorenna Mendes.

## Abstract

Developmental psychopathology has been a fruitful space for the investigation of the validity of multidimensionality. Nonetheless, previous studies in the field have not tested a model with a fine-grained division within the internalizing and externalizing spectra that also accounts for the contribution of a general psychopathology factor. The aim of this study is to test the validity and reliability of a theoretically-driven bifactor model with one general dimension and ten specific dimensions of psychopathology: *fear, somatic complaints, distressful thoughts, low mood, low motivation*/*energy, inattention-hyperactivity, temper loss, aggression, noncompliance*, and *low concern for others*. Our study evaluated 2,511 children aged 6-14 years-old from the Brazilian High-Risk Cohort for Psychiatric Disorders. There was support for the model in terms of its fit to the data, reliability, and external validity. Our findings highlight the potential of combined fine-grained approaches with models disentangling unity from diversity in developmental psychopathology.

## INTRODUCTION

Over development, children and adolescents may present a myriad of emotional and behavioral symptoms qualified as deviant or abnormal. At one hand, there is “diversity”, the multidimensional and specific quality that symptoms present, resulting in distinguishable syndromes. At the other hand, there is “unity”, the global quality that symptoms share with each other, resulting in a common cluster encompassing variance from symptoms of different clusters. In this study, we aimed to test the reliability and validity of a theoretically-driven model based on dimensions of developmental psychopathology using a bifactor structure to disentangle the roles of diversity and unity in psychopathological dimensions. Advancing our understanding of diversity and unity may lead to the identification of valid sub-types of existing disorders[1] and to the elucidation of divergent clinical implications of comorbidity[2].

Developmental psychopathology has been a fruitful space for the examination of the validity of *diversity*. Validity is the degree to which a diagnosis from a test correlates with expected external validators [3]. For instance, validity of distinctions among dimensions from the externalizing spectrum has been widely shown. The Oppositional Defiant Disorder (ODD) dimensions — irritable, headstrong, and hurtful have been associated with different genetic influences [4] and different outcomes both in children [5] and in adults [6]. The presence or absence of callous/unemotional traits in Conduct Disorder (CD) diagnosis has been linked to different brain processing mechanisms [7,8], distinct longitudinal outcomes [8,9], and different responses to treatment [7]. In contrast, distinctions among inattention, hyperactivity and impulsivity – dimensions from Attention-Deficit/Hyperactivity Disorder (ADHD) – have failed to provide valuable predictive information [10,11].

There is also evidence for valid distinctions among internalizing disorders. Fear and distress [12]–[14] have distinct genetic influences [15], different temperamental origins [16], divergent threat-related attentional biases[17], and contrasting patterns of activation in executive networks as measured by functional magnetic resonance imaging (MRI) [18]. Specificity also exists in dimensions of depression such as low mood and low motivation/energy as presented by anhedonia, including differences in treatment response[19], and in functional MRI features [20]. Moreover, it is also important to differentiate the somatic complaints cluster from other dimensions, since it is best conceptualized as separate from other aspects of psychopathology [21].

Nevertheless, despite abundant evidence on the differential validity of distinct dimensions of psychopathology, scores capturing variability on those dimensions tend to be highly inter-correlated. Therefore, growing attention has been directed to the validity of *unity* across all symptoms. These high rates of co-occurrence of psychiatric symptoms and disorders have led researchers to propose the existence of an overarching factor in psychopathology that captures common susceptibility to all psychiatric syndromes[22]. Caspi and colleagues, [23] coined the term “p-factor” to characterize this general psychopathology factor.

Despite the recent proliferation of models aiming to separate general from specific aspects of psychopathology, there is an absence of models aiming to capture the distinctions of psychopathological dimensions within the broad externalizing and internalizing spectrum. As reviewed above, these distinctions do exist; however, they need to be investigated with a model also able to parse the contribution of the general psychopathology dimension. Therefore, we proposed a model whose dimensions have shown differential validity in previous studies from developmental psychopathology.

The current study uses dimensions from the disruptive behavior model proposed by Wakschlag et al., [24]–[27] which conceptualizes four dimensions: *temper loss*, *noncompliance*, *aggression*, and *low concern for others*. We added to the disruptive behavior model, the following six dimensions: *fear, somatic complaints, distressful thoughts, low mood, low motivation*-*energy and inattention-hyperactivity*.

Therefore, the proposed model investigates 10 dimensions of developmental psychopathology, as follows:

a. *fear*: reflecting a tendency to react with fear and avoidance to unfamiliar stimuli, real or perceived threat [28, 29];
b. *distressful thoughts*: reflecting a tendency to react with rumination and preoccupation, accompanied by a sense of uncontrollability, to stressful situations including intensity, frequency and modulation [12], [30];
c. *somatic complaints*: reflecting a tendency to react to stressful situations with physical symptoms and heightened focus on visceral/bodily sensations [31];
d. *low mood*: reflecting persistent sadness, guilty and low self-esteem involving difficulties in being able to appropriately resolve negative emotions [32]–[34];
e. *low energy-motivation*: reflecting a tendency to present diminished motivation, energy or desire to engage in age-appropriate activities that are normally reinforcing [20], [34], [35];
f. *inattention-hyperactivity*: reflecting a tendency to present inattention, distractibility, lack of persistence, disorganization, excessive motor activity and impulsiveness DSM-5 [36]
g. *temper loss*: reflecting problems in regulation of anger including temper tantrums and angry mood [25], [27], [37];
h. *aggression:* reflecting a tendency to respond aggressively including multiple triggers and targets [25], [27];
i. *noncompliance*: reflecting resistance to comply with rules, and social norms [25], [27]; and
j. *low concern for others*: that parallels the callous unemotional dimension observed by Frick &White [8] reflecting a lack of empathy[8], [27].

Following the work from Robins and Guze [38], we aimed to provide validity for the proposed dimensions using three key validators. “Clinical description”, relying on differences related to sex/gender as a clinical feature, where externalizing dimensions are more prevalent among boys and internalizing dimensions more prevalent among girls[23], [39], [40]. “Laboratory studies”, by which we focus on executive function deficits for its relatedness with poor use of adaptive emotion regulation strategies [41], a characteristic we hypothesize to be a trans-diagnostic intermediate phenotype for both emotional and behavioral disorders [42]. Lastly, we rely on “delimitation from other disorders”, by using differences on dimensional scores among clinical groups of non-comorbid (pure) psychiatric disorders: phobias (e.g. separation anxiety, specific phobia), distress disorders (e.g., generalized anxiety, major depression), attention-deficit/hyperactivity disorder, and oppositional defiant/conduct disorders.

Here we use data from a large community sample of 6 to 14 year-old children [43] using a bifactor model to separate unity from diversity of ten psychopathological dimensions while also exploring external correlates. Moreover, to further develop previous evidence, we also used bifactor model-based psychometric indices to evaluate reliability of studied dimensions. We hypothesized that: a) The bifactor model with ten specific dimensions: *fear, somatic complaints, distress, low mood, low energy*/*motivation, inattention-hyperactivity, temper loss, noncompliance, aggression*, and *low concern for others* and a general psychopathology factor would provide a better fit to the data when compared to 5 competing models; b) The bifactor model-based psychometric indices of reliability would indicate that general and specific dimensions are reliable; c) external validators would support the existence of meaningful differences among the constructed general and specific dimensions.

## METHODS

### Sample and study design

Participants were part of the “Brazilian High-Risk Cohort Study for Psychiatric Disorders” [43] a large school based community study which includes a total of 57 state maintained schools (22 in Porto Alegre and 35 in São Paulo- Brazil). First, a screening phase was performed with families at public schools located close to research centers in Porto Alegre and São Paulo in the school registry day. After the families’ screening phase (8,012 families), we recruited two subgroups: one randomly selected (*n*=958) and one constituting a high-risk sample (*n*=1,553). The total sample was a combination of children from those two subgroups *(n*=*2,511)*. The children were aged 6 to 14 years-old (mean age = 9.65 years; SD=1.93) and 46.2% of them were girls. The high-risk sample selection involved a risk-prioritization procedure conducted targeting identification of individuals with current symptoms and/or a family history of specific disorders. Detailed information about the selection procedure can be found in [43]. The ethics committee of the University of São Paulo approved the study, and written consent was obtained from parents of all participants.

### Procedures

Assessments were: a) a household interview with parents conducted by a lay interviewer with the main caregiver which included measures of psychopathology such as the Children Behavior Checklist (CBCL)[44]; the Strengths and Difficulties Questionnaire (SDQ)[45]; and the Development and Well-Being Assessment (DAWBA)[46] and; b) child neurocognitive testing performed by trained psychologists and speech therapists at the school or home that included tasks detailed in the next section evaluating working memory, inhibitory control and temporal processing.

### Instruments

#### Child psychopathology

*CBCL* [44]— The Children Behavior Checklist CBCL is a rating scale composed by 112 items covering behavioral or emotional problems that have occurred during the past 6 months scored on a three-point scale: 0 (not true), 1 (somewhat or sometimes true), and 2 (very true or often.). The parent rated version was used in this study.

*SDQ* — The Strength and Difficulties Questionnaire [45] is a rating scale composed by 25 items evaluating emotional symptoms, conduct problems, hyperactivity, and peer problems that are scored on a three point scale: 0 (not true); 1 (somewhat true) and 2 (certainly true). The parent and teacher rated versions were used in this study.

*Construction of the 10-dimension model of psychopathology* — We constructed a model focusing on dimensions of psychopathology previously shown to have external and predictive validity. The ten proposed dimensions were: *fear, somatic complaints, distressful thoughts, low mood, low energy-motivation, temper loss, inattention-hyperactivity, noncompliance, aggression* and *low concern for others*. We determined, *a priori*, the number of dimensions to be equally distributed across the internalizing-externalizing division – the first five dimensions defined a priori as internalizing and the last five as externalizing. The purpose of this division was to produce a general factor equally balanced by externalizing and internalizing items in order to have a stable parameter estimation as advised by previous researchers [47]. From the item pool of 112 CBCL items and 25 SDQ items, we choose items which would represent our theoretically defined dimensions. After that, we iteratively performed Confirmatory Factor Analysis using bifactor models to select items presenting higher factor loadings both for general and specific dimensions. Lastly, we balanced the number of items for each dimensions (four items each) aiming the construction of a balanced (externalizing-internalizing) p-factor [23]. The resulting model was formed by 40 items (29 from CBCL and 11 from SDQ). See table 1.

**Table 1A.** Standardized factor loadings of the internalizing dimensions from the Non-orthogonal bifactor model.

| Factor | Item description | Factor loadings |  |
| --- | --- | --- | --- |
|  |  | G | S |
| **FEAR** | “Clings to adults or too dependent”[44] | 0.56 | 0.31 |
|  | “Fears certain animals, situations or places other than school”[44] | 0.26 | 0.48 |
|  | “Too shy or timid”[44] | 0.34 | 0.36 |
|  | “Many fears easily scared”[45] | 0.28 | 0.68 |
| **SOMA** | “Physical problems without known physical cause - Aches or pains”[44] | 0.46 | 0.52 |
|  | “Physical problems without known physical cause – nausea, feels sick”[44] | 0.34 | 0.63 |
|  | “Often complains of headaches, stomach-aches or sickness”[44] | 0.43 | 0.63 |
|  | “Feels dizzy or lightheaded”[45] | 0.30 | 0.67 |
| **DIST** | “Worries”[44] | 0.72 | 0.28 |
|  | “Many worries or often seems worried”[44] | 0.36 | 0.27 |
|  | “Can’t get his/her mind from certain thoughts/obsessions” [44] | 0.41 | 0.59 |
|  | “Fears he/she might think or do something bad” [45] | 0.17 | 0.71 |
| **LMOO** | “Unhappy, sad, or depressed” [44] | 0.74 | 0.33 |
|  | “Often unhappy, depressed or tearful”[44] | 0.68 | 0.45 |
|  | “Feels worthless or inferior”[44] | 0.49 | 0.58 |
|  | “Feels or complains that nobody loves him/her”[45] | 0.30 | 0.78 |
| **LOWM** | “There is very little he/she enjoys”[44] | 0.69 | 0.37 |
|  | “Underactive, slow moving or lacks energy” [44] | 0.47 | 0.53 |
|  | “Overtired without good reason”[44] | 0.65 | 0.45 |
|  | “Would rather be alone, than with others”[44] | 0.41 | 0.56 |

Abbreviations: G= general; S= specific; SOMA= somatic complaints; DIST= distress; LMOO= low mood; LOWM= anhedonia/low energy; IN/HYP= Inattention/hyperactivity; TEMP= temper loss; NONC= noncompliance; AGGR= aggression; LOWC= Low concern for others.

**Table 1B.** Standardized factor loadings of the externalizing dimensions from the Non-orthogonal bifactor model.

| Factor | Item description | Factor loadings |  |
| --- | --- | --- | --- |
|  |  | G | S |
| **IN/HYP** | “Inattentive or easily distracted”[44] | 0.63 | 0.52 |
|  | “Can’t concentrate, can’t pay attention for long”[44] | 0.60 | 0.63 |
|  | “Restless, cannot sit still for long”[45] | 0.34 | 0.68 |
|  | “Can’t sit still, restless or, hyperactive”[44] | 0.48 | 0.72 |
| **TEMP** | “Temper tantrums or hot temper”[44] | 0.65 | 0.50 |
|  | “Sudden changes in mood or feelings”[44] | 0.75 | 0.41 |
|  | “Sulks a lot”[44] | 0.75 | 0.40 |
|  | “Often loses temper”[45] | 0.31 | 0.85 |
| **AGGR** | “Physically attacks people”[44] | 0.42 | 0.69 |
|  | “Gets in many fights”[44] | 0.48 | 0.72 |
|  | “Threatens people”[44] | 0.49 | 0.59 |
|  | “Often fights with other youth or bullies them”[45] | 0.34 | 0.67 |
| **NONC** | “Disobedient at home”[44] | 0.59 | 0.64 |
|  | “Disobedient at school”[44] | 0.48 | 0.70 |
|  | “Breaks rules at home, at school, or elsewhere”[44] | 0.64 | 0.61 |
|  | “Generally well behaved, do what adults request*”[45] | 0.32 | 0.56 |
| **LOWC** | “Doesn’t seem to feel guilty after misbehaving”[44] | 0.63 | 0.38 |
|  | “Often offers to help others (parents, teachers, children)”[45] | 0.12 | 0.49 |
|  | “Considerate of other people’s feelings*”[45] | 0.30 | 0.71 |
|  | “Helpful if someone is hurt, upset or feeling ill*”[45] | 0.06 | 0.62 |

Abbreviations: G= general; S= specific; SOMA= somatic complaints; DIST= distress; LMOO= low mood; LOWM= anhedonia/low energy; IN/HYP= Inattention/hyperactivity; TEMP= temper loss; NONC= noncompliance; AGGR= aggression; LOWC= Low concern for others. \* Reverse scored.

*Psychiatric diagnosis*: Psychiatric disorders were diagnosed using the Development and Well-Being Assessment (DAWBA)[46], a structured interview that follows DSM-IV[48] criteria and evaluates child mental health symptoms. A team of nine psychiatrists supervised by a senior child psychiatrist rated data from DAWBA interviews that were administered to biological parents in accordance with previously reported procedures [46]. Based on DSM-IV, we selected six non-overlapping groups (e.g., no comorbidity allowed, pure psychiatric diagnosis) to be used as external validators which were; (1) *Typical developing controls* (TDC, n=1911) – absence of any psychiatric disorder; (2) *Phobic disorders* (n=112): separation anxiety (n=42), specific phobia (n=61), social phobia (n=18), agoraphobia (=3); (3) *Distress disorders* (n=94): generalized anxiety (n=30), major depression (n=35), not-otherwise specified depression (n=5), obsessive compulsive (n=5), or post-traumatic stress disorder (n=9); and (4) *ADHD* (n=146) – combined (n=9), inattentive (n=61), hyperactive/impulsive (n=25), not-otherwise specified (n=21); and (5) *ODD*/*CD (n*=*60)* – ODD (n=44), CD (n=14) and not-otherwise specified (n=5).

#### Child Cognition

Child cognitive data was available for at least one measure for 2,395 children. Descriptive statistics for each of the measures can be found in supplemental material (Supplemental Table S1).

##### Working memory

a) Digit span: This task is a sub-set of WISC-III [49] that evaluates verbal working memory by asking participants to repeat sequences of numbers to whom they were orally presented for either as heard (forward version) or in reverse order (backwards version) with increasing level of difficulty. The dependent measure was the level at which the participant failed to correctly repeat the numbers on two consecutive trials at one level of difficulty.

b) Corsi blocks task [50]: This task evaluates the visuo-spatial working memory by asking participants to observe the researcher as he/she taps a sequence of up to nine identical spatially separated blocks and then mimic the pattern presented by the researcher with increasing level of difficulty in sequential trials. The dependent measure was the level at which the participant failed to correctly repeat the sequence of blocks on two consecutive trials at one level of difficulty.

##### Inhibitory-control

c) Conflict Control Task [51]: This task evaluates the executive component of inhibitory control by creating a dominant motor response and then asking participants to occasionally suppress this preponderant tendency and initiate a motor action in the opposite direction. First, participants are instructed to press the button indicating the arrow direction appearing in the computer screen (congruent trials) in a total of 75 trials when the arrow color was green. The remaining 25 incongruent trials were presented with red arrows in which participants had to respond in the opposite direction to that indicated by the arrows (a conflict effect). Inter-trial interval was 1500 msec and the stimulus duration was 100 msec. Accuracy and speed was equally emphasized in task instructions. The dependent measure was the percentage of correct responses in the incongruent trials.

d) Go/No-Go [52]: This task assessed the participants capacity to completely suppress a preponderant tendency to press the button indicating the direction of the green arrows (Go stimuli, n= 75) when a double-headed green arrow (No-Go stimuli; n=25) appeared in the screen. This task consisted of 100 trials with stimulus duration of 100 msec and inter-trial interval of 1500 msec. Accuracy and speed was equally emphasized in task instructions. The dependent measure was the percentage of failed inhibitions in the no-go trials (commission errors).

##### Time processing

e) Time Anticipation tasks – 400ms and 2000ms (TA;[53];[54]): This game-like task examined the ability to modulate the time of a motor response to precisely coincide with the onset of a visual stimulus. A brief narrative was created to help participants understand the task: they were told they were space explorers that had to refurnish an allied spaceship running out of oxygen in order to save the allied crew by anticipating when the target (spaceship) would reappear. In each task, the allied spaceship was visible for the first 10 trials, and for the remaining 16 trials participants were asked to press a button to anticipate when it would arrive, because an invisible shield was activated. Participants had a 750 msec window of time to respond correctly and received feedback after every trial. In task 1, the anticipation interval was 400 milliseconds and in task 2 it was 2,000 milliseconds. The 2000 ms task was always administered after the 400ms task. The dependent measure for both the 400ms and 2000ms anticipation tasks was the mean percentage of total hits (button pressed in correct time window interval).

##### Child Executive Function Model

We used a second-order model previously validated [55] developed to resume child cognitive data that encompassed a second-order executive function Factor (EF) and three lower-order factors (i.e., Working Memory [WM]; Inhibitory Control [IC]; and Temporal Processing [TP]) loading onto the superordinate EF factor. As previously found [55], this model exhibited excellent fit to the data (RMSEA=0.004, CI90% <0.001-0.027, CFI>0.999, TLI=0.999) and its lower-order factors exhibited high loadings on the higher-order factor (λ ranging from 0.4 to 0.8).

### Data Analysis

*First*, we examined the structure of psychopathology testing six models (table2): a) One factor model; b) Correlated Model with two dimensions: internalizing and externalizing; c) Correlated Model with ten specific dimensions; d) Non-orthogonal Bifactor Model with one general factor and two specific dimensions; e) Orthogonal Bifactor Model with one general factor and ten specific dimensions and; f) Non-orthogonal Bifactor Model with one general factor and ten specific factors allowed to correlate between each other. Models were compared using the following indexes: Chi-square fit statistics, the Comparative Fit Index (CFI), the Tucker Lewis Index (TLI), and the Root Mean Square Error of Approximation (RMSEA) as recommended by [56], [57]. CFI and TLI above 0.95 and RMSEA below 0.06 indicate good model fit [56].

**Table 2.** Model fit indices for the 6 tested models of psychopathology

| Fit indices | One factor model | Correlated Model (2 factors: int x ext) | Correlated Model (10 factors) | Non-Orthogonal Bifactor Model (1 general factor and 2 specific factors; int x ext) | Orthogonal Bifactor Model* (1 general factor and 10 specific factors) | Non-Orthogonal Bifactor Model (1 general factor and 10 specific factors) |
| --- | --- | --- | --- | --- | --- | --- |
| $\chi^2$ | 173210.515(df=780; p<0.001) | 173210.515(df=780;p<0.001) | 173210.515(df=780; p<0.001) | 173210.515(df=780;p<0.001) | * Matrix psi non positive definite. | 173210.515(df =780; p<0.001) |
| CFI | 0.947 | 0.964 | 0.984 | 0.982 |  | 0.992 |
| TLI | 0.944 | 0.962 | 0.982 | 0.980 |  | 0.991 |
| RMSEA | 0.070 | 0.058 | 0.040 | 0.042 |  | 0.029 |
| CI 90% RMSEA | (0.069-0.071) | (0.056-0.059) | (0.039-0.041) | (0.04-0.043) |  | (0.027-0.03) |
| WRMR | 3.389 | 2.832 | 2.007 | 2.086 |  | 1.532 |

Note: Int= internalizing; Ext= externalizing; $\chi^2$ = Chi Square and; df= degrees of freedom; CFI= Comparative Fit Index; TLI= Tucker-Lewis Index; RMSEA= Root-Mean-Square Error of Approximation; CI= Confidence Interval; and WRMSR= Weighted Root Mean Square Residual.

*Second*, we tested the reliability (table3) considering the following coefficients that may vary between 0 and 1, where higher scores indicate greater reliability: (a) Omega (ω); [58]–[61] and (b) Omega subscale (ωs) [59], [60]; both of them assess the systematic variance affecting unit-weighted composite scores attributable to multiple common factors, where higher values indicate a reliable multidimensional composite; (c) Omega Hierarchical (ωH);[58]–[60], [62] assesses the degree to which composite scale scores are interpretable as measure of a single common factor that considers the error variance; (d) Omega hierarchical subscale (ωhs); [59], [60] measures the reliability of subscale scores after removing the effects of the general factor.

*Lastly*, we evaluated the validity of the dimensions by using executive function measures and sex as external validators in structural equation models. In addition, we performed a series of Analysis of Variance (ANOVA) using psychopathological dimensions scores as the dependent measures and clinical non-overlapping groups as independent variables.

Analyses were performed in R software program, version 3.2.3 [63] with lavaan package [64] using Full Information Maximum Likelihood (FIML) estimation.

## RESULTS

### Structure of psychopathology

The non-orthogonal bifactor model with a general psychopathology factor (g) and ten specific factors: *fear, somatic complaints, distressful thoughts, low mood, low energy-motivation, temper loss, inattention-hyperactivity, noncompliance, aggression and low concern for others* exhibited the best fit to the data when compared to the other five models: CFI= 0.992, TLI= 0.991, RMSEA= 0.029, 90%, and confidence interval (CI) = (0.027 – 0.03) (table 2).

On figure 1 the latent correlations of the non-orthogonal bifactor model (the model accounting for the general factor) are shown. The non-orthogonal bifactor model is able to elucidate which correlations between dimensions are not statistically significant after partitioning the variance attributed to the general psychopathology factor; and also to show expected negative correlations between dimensions (*distressful thoughts* and *noncompliance*; *distressful thoughts* and *low concern for others*). Of note, despite *temper loss* traditional classification as an externalizing dimension, this factor showed moderate positive correlations with most dimensions considered internalizing.

**Figure 1.**
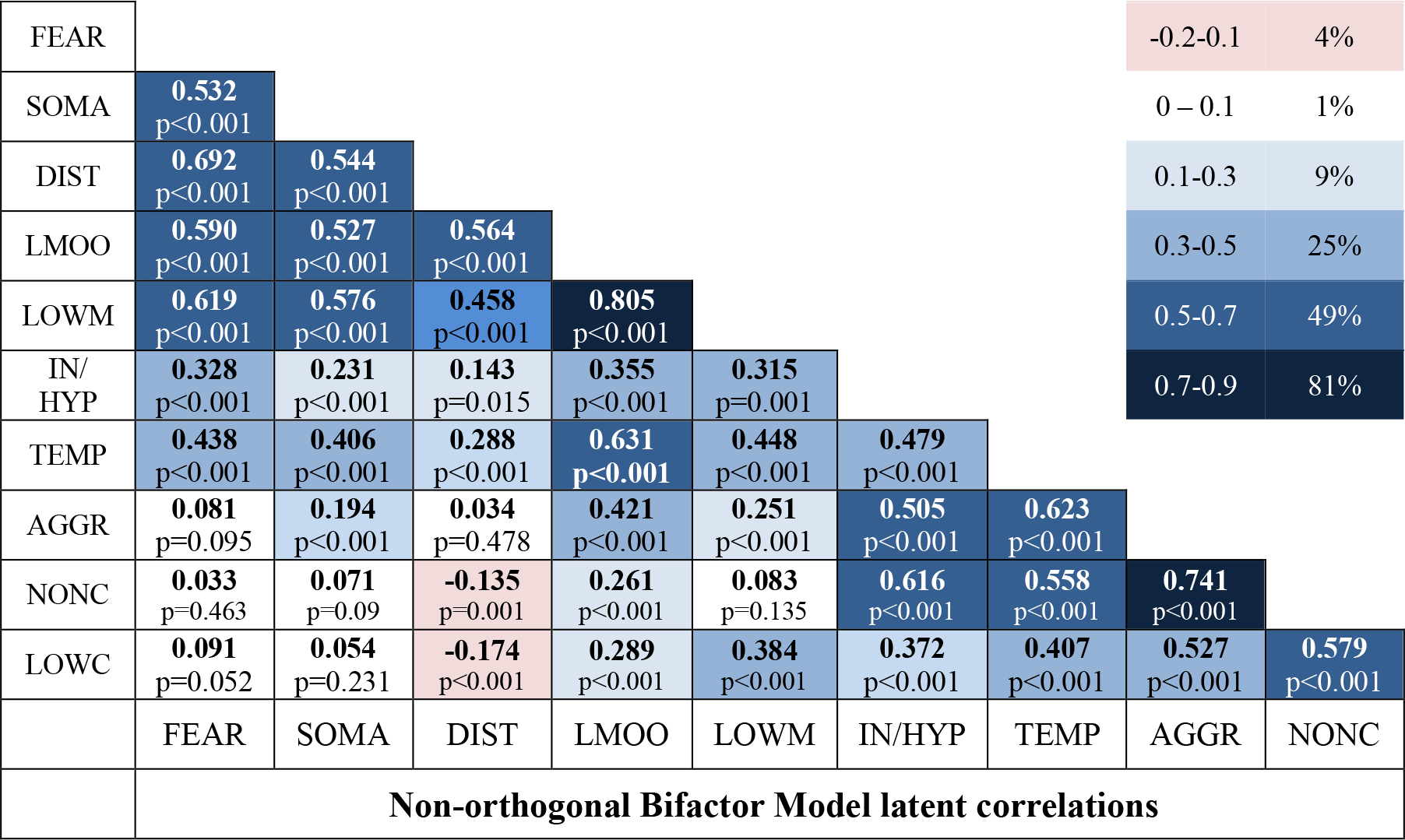
Latent Correlations among psychopathological dimensions from the Non Orthogonal Bifactor model. On the top right, a legend displays the colors for each correlation level between dimensions. Note: SOMA= *somatic complaints*; DIST= *distressful thoughts*; LMOO= *low mood*; LOWM= *low motivation*/*energy*; IN/HYP= *Inattention*/*hyperactivity*; TEMP= *temper loss*; NONC= *noncompliance*; and LOWC= *low concern for others*.

### Reliability

As shown in table 3, the omega for the general factor was high *ω= 0.94)* as was the omega subscale values *(ω*_*s*_ = *0.94)* indicating a highly reliable multidimensional composite. The hierarchical omega for the general factor was *0.82*, meaning that *82%* of the variance of unit-weighted total scores can be attributed to the individual differences on the general factor. When comparing the *ω*_*h*_ *(0.82)* with *ω (0.94)*, one finds that most of the reliable variance in total scores *87% (0.82/0.94*= *0.87)* are due to individual differences in the general factor; that only *12% (0.94-0.82*= *0.12)* of the reliable variance of the total scores can be attributed to the multidimensionality caused by the group factors and that *6% (1-0.94)* of the variance is due to random error. The small values of *ω*_*hs*_ *(ranging from 0.01 to 0.02)* suggest that the assumed reliability of *ω*s was due to individual differences on the general factor.

**Table 3.** Reliability indices from the non-orthogonal bifactor model of psychopathology with ten dimensions.

| Dimensions | Reliability indices |  |  |  |
| --- | --- | --- | --- | --- |
| | Omega ($\omega$) | Omega Hierarchical ($\omega_H$) | Omega Subscale ($\omega_s$) | Omega hierarchical subscale ($\omega_{hs}$) |
| General Psychopathology | 0.94 | 0.82 | - | - |
| Fear | - | - | 0.94 | 0.01 |
| Somatic Complaints | - | - | 0.94 | 0.02 |
| Distressful thoughts | - | - | 0.94 | 0.01 |
| Low Mood | - | - | 0.94 | 0.01 |
| Low motivation/energy | - | - | 0.94 | 0.01 |
| Inattention/hyperactivity | - | - | 0.94 | 0.02 |
| Temper Loss | - | - | 0.94 | 0.01 |
| Aggression | - | - | 0.94 | 0.02 |
| Noncompliance | - | - | 0.94 | 0.02 |
| Low Concern for others | - | - | 0.94 | 0.01 |

### External correlates

#### Clinical description: sex/gender differences

We found a clear sex/gender effect (figure 2) after separating general psychopathology from specific dimensions: boys presented higher scores on externalizing dimensions (*inattention-hyperactivity, noncompliance and aggression*), whereas girls presented higher scores on most internalizing dimensions (*fear*, *somatic complaints*, *distressful thoughts* and *low mood*). The general psychopathology factor, *low motivation/energy, temper loss*, and *low concern for others* were not associated with sex/gender.

**Figure 2.**
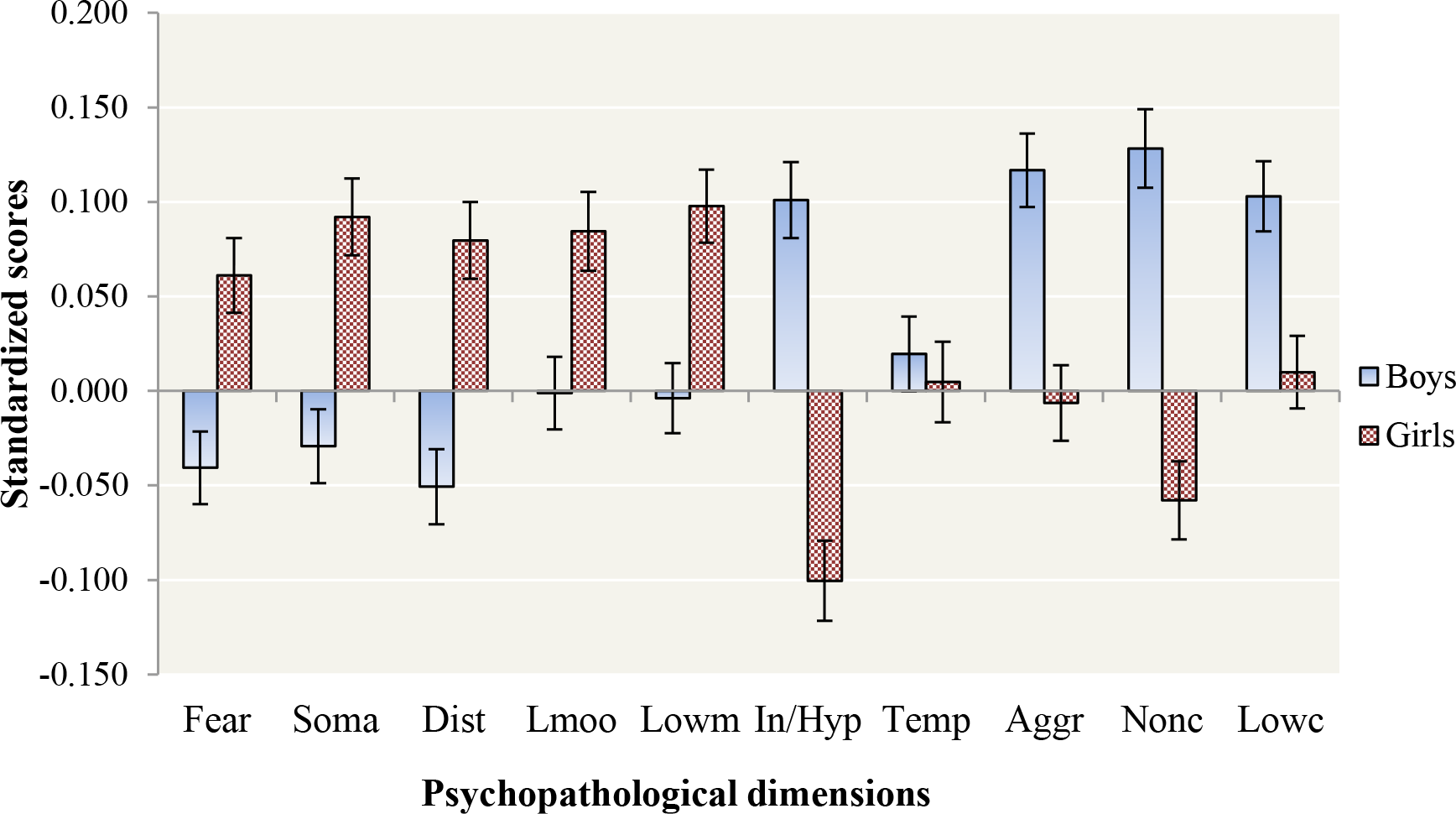
Sex/gender differences on psychopathological dimensions. Abbreviations: SOMA= somatic complaints; DIST= distressful thoughts; LMOO= low mood; LOWM= low motivation/energy; IN/HYP= Inattention/hyperactivity; TEMP= temper loss; NONC= noncompliance; and LOWC= low concern for others.

#### Laboratory tests: executive function dimensions and psychopathological dimensions

Regressions of child psychopathological dimensions on global executive function (Figure 3) revealed negative associations between global executive function and *general psychopathology (β*= −*0.15, p*<*0.001)*, *fear (β*=−*0.138, p*=*0.001), and inattention-hyperactivity (β*=−*0.144, p*<*0.001)*. On the other hand, better global executive function predicted more *distressful thoughts (β*=*0.137, p*<*0.001) and low mood (β*=*0.077, p*=*0.034)*. No associations were found for *somatic complaints (β*= *0.062, p*=*0.077), low motivation/energy (β*=*0.053, p*=*0.144), temper loss (β*=*0.027, p*=*0.485), aggression (β*=−*0.048, p*=*0.217), noncompliance (β*=*0.014, p*=*0.720), and low concern for others (β*=*0.120, p*=*0.290)*.

**Figure 3.**
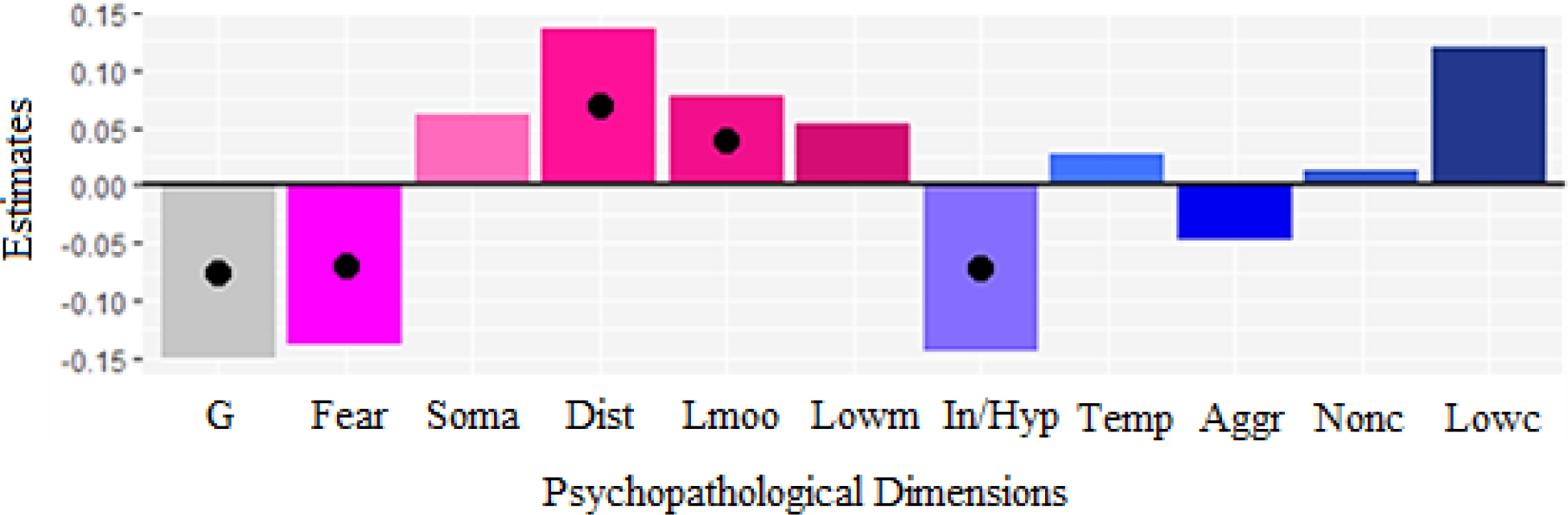
Regression Coefficients (β) of psychopathological dimensions on global executive function. Abbreviations: G= General psychopathology; Soma= somatic complaints; Dist= distressful thoughts; Lmoo= low mood; Lowm= low motivation/energy; In/hyp= Inattention/hyperactivity; Temp= temper loss; Nonc= noncompliance; and Lowc= low concern for others. Note: (*) represents 0.05 level of significance.

#### Delimitation from clinical disorders: clinical associations

ANOVAs demonstrated an effect of group (figure 4) on all psychopathological dimensions. As depicted in figure 4; TDC group presented the lowest standardized scores on psychopathological dimensions. The phobias group tended to show significantly less noncompliance and low concern for others than the TDC group. Phobias group also presented significantly lower scores on general psychopathology dimension when compared to distress group and to ODD/CD. Phobias and distress groups presented similar levels of *fear, somatic complaints* and *distressful thoughts* scores, however, the distress group had higher scores *on low motivation*/*energy* and *low mood*, and also displayed significantly higher scores on dimensions considered externalizing (*temper loss, aggression, low concern for others*) than the phobias group. As expected, ADHD group presented significantly less general psychopathology, temper loss, aggression and low concern for others than the ODD/CD group. When compared to the other groups, ODD/CD group showed significantly lower scores on *distressful thoughts* and *fear* dimensions.

**Figure 4.**
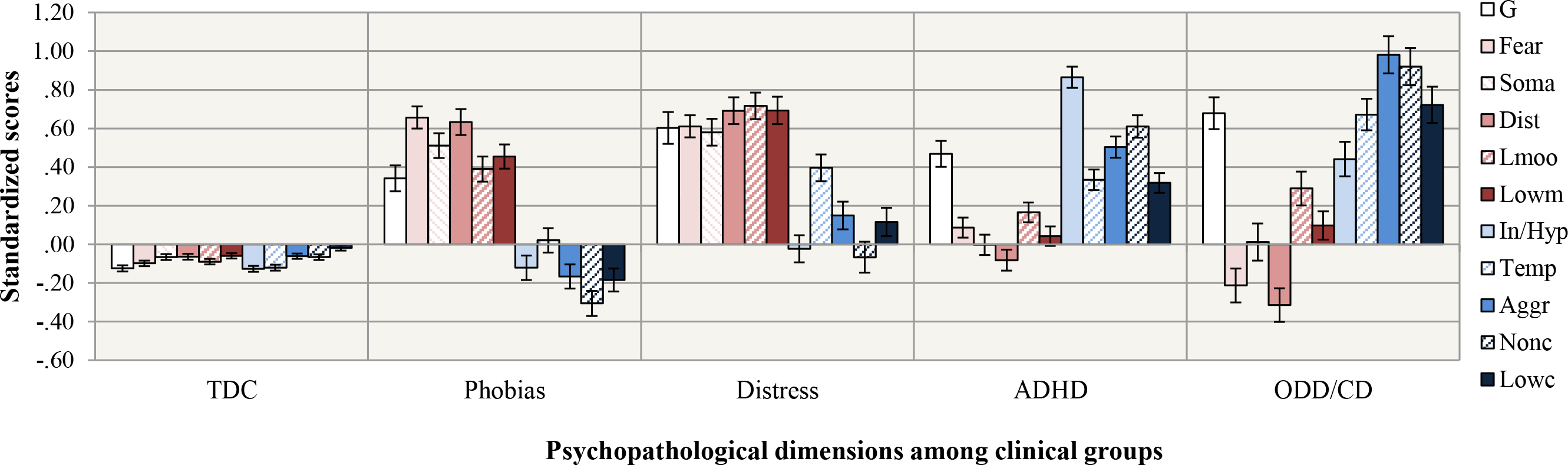
Comparison of between group differences on the standardized scores of psychopathological dimensions. Abbreviations: Soma= somatic complaints; Dist= distressful thoughts; Lmoo= low mood; Lowm= low motivation/energy; In/hyp= Inattention/hyperactivity; Temp= temper loss; Nonc= noncompliance; and Lowc= low concern for others; TDC, Typically Developing Children; ADHD, Attention Deficit/Hyperactivity Disorder; ODD/CD, Oppositional Defiant Disorder/Conduct Disorder

## DISCUSSION

Our data investigated the reliability and validity of a model based on developmental psychopathology dimensions specifically constructed to investigate the reliability and validity of ten empirically defined psychopathological dimensions (*diversity*) after accounting for the general factor (*unity*) using a bifactor model. Our main findings can be summarized as follows. *First*, our model demonstrated good fit to the data and the proposed bifactor hierarchical structured allowed an interesting scrutiny of the latent correlations among dimensions of psychopathology. *Second*, when specific reliability indexes were investigated, we showed that most of the reliable variance in total scores was due to individual differences in the general factor, whereas reliability of specific factors was largely dependent upon the general factor. *Third*, despite low reliability of specific factors after accounting for the general factor, the proposed model was able to show differential validity of specific factors over and above the general factor in terms of distinct associations with sex/gender, global score of executive functions, and clinical indicators.

As described in other studies, the bifactor model presented the best goodness of fit as compared to other tested models. Differently from other studies, specific factors were allowed to correlate between each other (but not with the general factor). Since the proposed model has 10 specific dimensions, it is reasonable to assume that correlations between specific factors, which are not accounted only by the general factor, will emerge. In fact, the pattern of the residual correlations revealed interesting patterns. We found negative correlations between *low concern for others* and *distressful thoughts* and negative correlations between this last dimension and *noncompliance*. *Low concern for others* when paralleling the concept of callous unemotional traits [8] is associated with insensitivity to punishment [8], [65], [66] while the *distressful thoughts* dimension is associated with an enhanced detection of future punishment [67]. Therefore, one should expect to find at the latent level a negative correlation between them. Also, the compatibility previously found between rule compliance and normative moral emotions (distress following transgressions) [68] may explain our results on the negative correlation between *distressful thoughts* and *noncompliance*. Interestingly, the *temper loss dimension* had moderate correlations with dimensions from the externalizing spectrum but also presented significant correlations with dimensions from the internalizing spectrum; what is consistent with previous findings on the heterotypic comorbidity of irritability[69], [70] and heterotypic continuity of irritability predicting internalizing disorders later in life [5], [6], [71], [72].

In order to advance previous evidence we tested the reliability of general and specific dimensions with model-based psychometric indices [60]. We reported a high omega for the general factor, meaning that our total score was a highly reliable composite and that the general psychopathology factor was the most reliable source of variance of the model. Subscale scores also had reliable ω_s_. However, ω_H_ and ω_hs_ indicated that the reliability of specific factors was largely dependent upon the general factor. If one considers the general factor as a liability to deviate from normal to pathological development, this liability may be the most reliable construct available leaving the phenomenology of “how” this liability is presented – the specific factors – as more elusive constructs. Despite presenting low reliability, specific factors showed distinct associations with external validators, what provides support for their differential validity.

In terms of sex/gender associations, boys were more likely to present *inattention/hyperactivity*, *aggression* and *noncompliance*; whereas girls were more likely to present *fear*, *somatic complaints*, *distressful thoughts* and *low mood*. The female liability towards internalizing spectrum dimensions and the male liability towards the externalizing spectrum is consistent with findings from studies in adults [23] and youth [40] and might be an indicative of the role of sex/gender as one of the determinants of how general psychopathology factor will present phenomenologically.

The proposed model displayed differential validity in terms of executive function since we found a different profile for dimensions within the same broad spectrum. The proposed model confirmed previous findings of the association between worse executive function and the general psychopathology factor in youth [73], [55], [74], [75], [18] and the association between inattention-hyperactivity and worse executive function [11], [76], [77]. The bifactor model displayed an EF divergent profile for anxiety dimensions; worse performance related to *fear* and better performance related to *distressful thoughts*. These results contradict a previous finding on the negative association between anxiety and EF [78]. This difference can be due to a number of factors. *First*, differently from the previous work, we used a bifactor model structure that enabled us to investigate the association of specific psychopathology dimensions after accounting for the contribution of the *general psychopathology factor*. Second, the two dimensions linked to the anxiety construct (*distressful thoughts* and *fear*) were separated into different factors in our work. In addition, another study[18] that used a bifactor structure also found a tendency to better working memory performance for their anxious-misery dimension paralleling our *distressful thoughts dimension*. *Lastly*, we used a dimensional classification of psychopathology for a community sample, which is not expected to present severe symptomatology or impairment. In addition, better EF was also associated with higher scores on *low mood*. It is possible that *distressful thoughts* and *low mood* dimensions related to depressive symptoms are associated with more rumination [31], [79] a trait linked to better performance on a EF task requiring goal maintenance but more errors on a task requiring goal-shifting [80]. Therefore, the finding of this positive association between EF and these dimensions may be due to the absence of a task evaluating the EF shifting dimension in our study.

Our model provided good clinical concurrent validity since general and specific dimensions z scores were able to differentiate between the non-overlapping clinical groups. The distress group presented significantly more *low mood* than the phobias group what may mimic the finding of homotypic comorbidity between anxiety and depression [81] and the finding of childhood anxiety disorders homotypic continuity with adult depression being due to generalized anxiety disorder only [82]. Specific dimensions were also able to differentiate between ADHD group and ODD/CD group with the latter presenting more *low mood* than the ADHD group what is consistent with the ODD importance in mediating the relationship between ADHD and depression [5], [69].

The current work presents some limitations that should be noted. The model was not built with all items from CBCL or SDQ questionnaires. Indeed, only items specifically important to reflect the hypothesized dimensions were chosen to create the model. Moreover, we theorized all dimensions with the intention to mimic prior work on *fear*, *somatic complaints*, *distressful thoughts*, *low mood*, *low motivation/energy*, *inattention-hyperactivity* [11], [12], [20], [31] and dimensions from the developmental model of Wakschlag et al., [26]. However, the proposed dimensions have not been validated in relation to stablished measures formulated, what may impose measurement limitations. Nevertheless, despite being a new composite, the omega of the general and specific factors in our model were high, meaning it was a good multidimensional composite and the evaluation of dimensions from opposed spectra was also performed what could not be developed with validated measures from cited works. Therefore, replication of our findings with measures deliberately designed for this purpose is warranted; just as the creation of new instruments specially designed to separate general from specific contributions of child psychopathological dimensions. Nonetheless, this work has several strengths: it benefits from a large community sample; it evaluates psychopathology dimensionally, it evaluates the reliability of psychopathological dimensions with bifactor model psychometric indices and it probes dimensions specific neuropsychological deficits after accounting for the general psychopathology factor.

Our work supports the need to search for multidimensionality in developmental psychopathology and provides a testable theoretical framework for investigating and testing different validators from specific dimensions over and above the contribution of the general psychopathology factor. It is important for future works to evaluate if indeed the p factor is responsible for the transition from a normative developmental trajectory towards a trajectory of mental illness and to test with mixture models if the p factor may be responsible for the transition from normative misbehavior to full-blown mental illness.

## SUMMARY

Although there are many studies reporting the differential utility of multidimensionality in terms of external validators in developmental psychopathology, previous studies in the field have not tested a model with a fine-grained division within the internalizing and externalizing spectra while also accounting for the contribution of a general psychopathology factor. This study investigated the reliability and validity of a bifactor model with 1 general dimension (unity) and ten specific dimensions of psychopathology (diversity): *fear, somatic complaints, distressful thoughts, low mood, low motivation/energy, inattention-hyperactivity, temper loss, aggression, noncompliance*, and *low concern for others*. The proposed model was compared to five competing models and it provided a good fit to the data and it was supported by the reliability indexes. The bifactor model was able to show differential validity of specific factors over and above the general factor in terms of distinct associations with sex/gender, global score of executive function and clinical indicators. For instance, we found a different profile of Executive Function performance for dimensions within the same broad spectrum (Distressful thoughts × fear), (low mood × fear) and (inattention-hyperactivity × low concern for others) what justifies the creation of models with a more fine-grained division. This study provides a theoretical framework for investigating and testing differential validators of specific dimensions above and beyond the contribution of the general psychopathology factor.

