## Supplemental Table 1 for "Disentangling unity from diversity in developmental psychopathology"

### SUPPLEMENTAL MATERIAL

**Supplemental Table S1**- Descriptive statistics on executive function tasks (crude scores)

| | Mean | SD | p25 | p75 | Minimum | Maximum | Valid N |
|---|---|---|---|---|---|---|---|
| Digit span (backward) | 3.54 | 1.57 | 3.00 | 4.00 | 0 | 12.00 | 2243 |
| Corsi block (backward) | 4.80 | 2.10 | 3.00 | 6.00 | 0 | 14.00 | 2213 |
| CCT % Correct Inhibitions (inc) | .59 | .22 | .42 | .76 | 0 | 1 | 2166 |
| Go/No-Go: Comission | .25 | .22 | 0.08 | .38 | 0 | 1 | 2158 |
| Time Anticipation (0,4s): hits | .66 | .22 | .50 | .86 | 0 | 1 | 2185 |
| Time Anticipation (2s): hits | .33 | .25 | .143 | .500 | 0 | 1 | 2180 |
